# GMP-grade neural progenitor derivation and differentiation from clinical-grade human embryonic stem cells

**DOI:** 10.1101/2020.04.22.043216

**Authors:** Loriana Vitillo, Catherine Durance, Zoe Hewitt, Harry Moore, Austin Smith, Ludovic Vallier

**Affiliations:** Wellcome -MRC Cambridge Stem Cell Institute, Jeffrey Cheah Biomedical Centre, University of Cambridge, Cambridge, CB2 0AW, UK; The Centre for Stem Cell Biology, Alfred Denny Building, The University of Sheffield, Western Bank, Sheffield, S10 2TN, UK

## Abstract

A major challenge for the clinical use of human pluripotent stem cells is the development of safe, robust and controlled differentiation protocols. Adaptation of research protocols using reagents designated as research-only to those which are suitable for clinical use, often referred to as Good Manufacturing Practice (GMP) reagents, is a crucial and laborious step in the translational pipeline. However, published protocols to assist this process remain very limited. Here we present a new GMP-compliant protocol to derive long-term neuroepithelial stem cell progenitors (lt-NES), which are multipotent, bankable, and karyotypically stable. This protocol resulted in robust and reproducible differentiation of several clinical-grade embryonic stem cells, deposited in the UK Stem Cell Bank, from which we derived lt-NES. Furthermore, GMP-derived lt-NES demonstrated a high neurogenic potential while retaining the ability to be redirected to several neuronal sub-types. Overall, we report the feasibility of derivation and differentiation of clinical grade embryonic stem cell lines into lt-NES under GMP-compliant conditions. Our protocols could be used as a flexible tool to speed up translation-to-clinic of pluripotent stem cells for a variety of neurological therapies or regenerative medicine studies.

## INTRODUCTION

The stem cell revolution started with the isolation of human embryonic stem cells (hESCs) (Thomson et al., 1998) followed by the arrival of induced pluripotent stem cells (iPSCs) (Takahashi and Yamakana, 2006), and their differentiation to an ever-increasing number of cell types, has led to the prospect of shifting medicine to a new paradigm based on cellular repair. Despite this enticing prospect, the number of clinical trials based on human pluripotent stem cells (hPSCs) remain limited when compared to other cell types (Trounson and McDonald, 2015; Ratcliffe et al., 2013). There is a consensus that hPSCs have a complex and distinct set of scientific, technical and regulatory bottlenecks that hamper their translation to clinical applications (Williams et al., 2016; Whiting et al., 2015; Ratcliffe et al., 2013; Abbasalizadeh and Baharvand, 2013).

One hurdle is that the often-large lists of raw materials used in differentiation protocols are designated of research-grade and were never intended for cell therapy applications. The developers of advanced therapeutic medicinal products (ATMPs) need to follow strict Good Manufacturing Practice (GMP) guidelines to ensure quality and safety of the end products before performing clinical trials. Therefore, raw materials used in differentiation protocols must meet these guidelines. Although policies around raw materials for cell therapeutics are currently flexible, clinical trials regulations require for each reagent to be extensively risk-assessed and qualified (Stacey et al., 2018). It is in this context that so called GMP-grade materials suitable for clinical use will facilitate clinical trial submission and will likely become standard in the field (Solomon et al., 2016).

To comply with such regulations, hESCs have been derived under clinical grade conditions (Hewitt et al., 2007; Jacquet et al., 2013; Wang et al., 2015) and GMP-compliant culture platforms have been developed (Chen et al., 2011; Nakagawa et al., 2014). However, published GMP-compliant differentiation protocols are still notably lacking, but their development would significantly speed the translational pipeline for pluripotent stem cells, particularly for academic groups.

Previous work, including ours, showed that hPSCs could be differentiated into a long-term neuroepithelial-like stem cells population, lt-NES, with stable neurogenic potential towards several neuronal subtypes (Koch et al. 2009; Falk et al., 2012). In the context of regenerative medicine, a source of neurons that is an expandable, bankable and intermediate (i.e. at progenitor stage) has several advantages over run-through protocols. Lt-NES would reduce processing steps, would be a convenient quality control check point and could potentially be used for several applications, facilitate scalability and also by-pass intrinsic line-to-line variability associated with iPSCs (Falk et al., 2012). Here, we develop a novel protocol for the derivation and differentiation of lt-NES from clinical-grade hESC lines deposited in the UK Stem Cell Bank based on GMP-grade media and factors.

## RESULTS

### Development of an efficient GMP-compatible protocol for lt-NES derivation

In order to develop a GMP-compatible protocol for lt-NES derivation, we used H9 hESCs routinely cultured on a widely recognized and defined culture system based on recombinant vitronectin and Essential 8 (E8) (Chen et al., 2011). We started by examining the performance of an embryoid body (EB)-based neural differentiation system under standard research grade (Falk et al., 2012) versus GMP media conditions. HESCs were allowed to aggregate spontaneously in suspension for 5 days in research-grade Knock-out Serum Replacement differentiation media (KSR) as previously described (Falk et al., 2012) or in GMP-grade Essential 6 (E6), which is composed similarly to GMP E8 but without bFGF or TGFβ and thus is suitable for differentiation. The hESCs formed compacted and round-shaped EBs in KSR while in E6 they showed an elevated level of attachment to the ultra-low adherence dish and disaggregated into smaller pieces over the 5-day period (Figure S1A). Consequently, we observed that poorly formed EBs in E6 were also inefficient during neural induction, assessed by the hallmark of neural rosettes, compared to those in KSR (Figure S1B).

Moreover, although neural differentiation is clearly possible via classic spontaneous EB formation methods, this system is not standardized, as the size and the shape of EBs is uncontrolled and this impacts on the reproducibility of differentiation and yield. Therefore, we next examined the performance of an alternative method to produce EBs of defined size based on seeding dissociated cells in microwells. HESCs were dissociated into single cells and seeded at a concentration of 10,000 hESCs per microwell in the presence of a GMP-grade ROCK inhibitor (Revitacell, Life Technologies) in either E6 or KSR. After 24hr, similar sized EBs were formed in both conditions and at day 5 they remained aggregated (Figure 1A and S1C). Upon dissociation, an equal number of same sized EBs were obtained with this protocol from both KSR and E6 media. We tested the neural induction efficiency of standardized EBs by looking for the emergence of neural rosettes after plating in neuronal inducing conditions. Surprisingly, neural rosettes emerged more prevalently from EBs derived from E6 rather than KSR, in contrast to the spontaneous differentiation system (Figure S1C). Moreover, GMP-neural induction was robust and highly efficient, as shown by rosettes forming simultaneously and similarly at the centre of plated EBs within 3 days (Movie 1). We also tested neural induction on a defined laminin matrix, laminin 521, now available in GMP-grade, showing similar results to standard research-grade poly-L-Ornithine/laminin substrate (Figure S1D).

**Figure 1.**
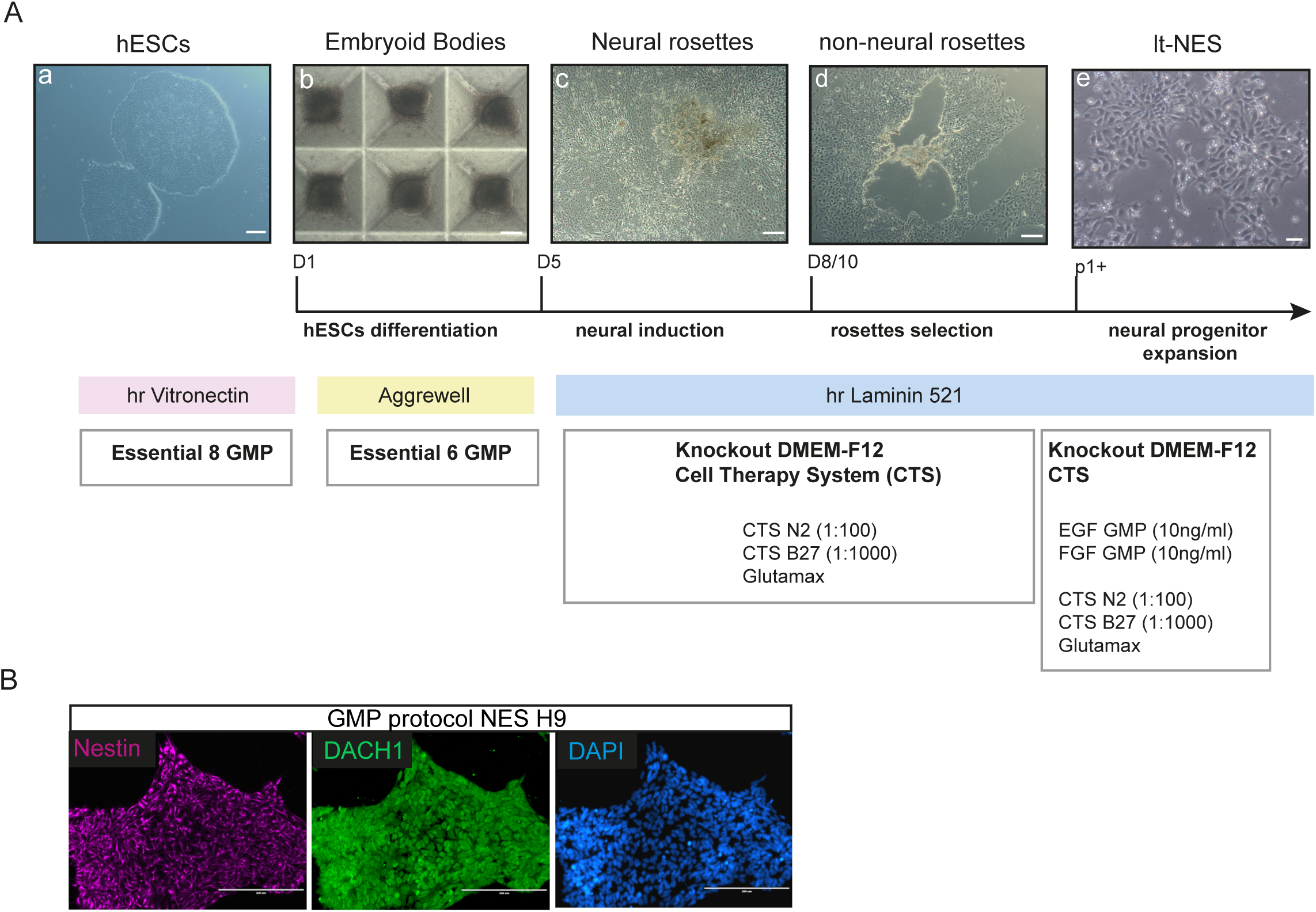
Development of an efficient GMP-compatible protocol for lt-NES. A) Step-by-step diagram of GMP-compatible differentiation protocol of hESCs into lt-NES. Scale bars a,c,d 100 μm; b 50 μm and e 20 μm. B) Immunofluorescence for lt-NES markers Nestin and Dach1 in research-grade line H9 after derivation with GMP-compatible protocol. Scale bars 200 μm.

Derivation of lt-NES from neural rosettes has previously been described using a manual picking system with needle dissection of rosettes (Koch et al., 2009; Falk et al., 2012). In the context of future manufacturing applications for lt-NES, we tested the suitability of a commercially available rosette selection solution (STEMdiff™ Neural Rosette Selection, Stem Cell Technologies) with our GMP protocol. Application of the reagent allowed the pipetting out of loosened neural rosette cluster from a surrounding non-neural rosette ring of differentiated cells without the need for manual dissection (Figure 1A). With this method, lt-NES were derived after disaggregating the rosettes and plating the cells at high density in presence of GMP-grade EGF and FGF, and cultured thereafter until sub-confluent. Under these culture conditions, it-NES could be expanded every 2-3 days, a result of their typical self-renewing capacity, while maintaining a highly pure population (Figure 1A). Indeed, characteristic lt-NES markers Nestin and Dach1 were expressed homogeneously by the cultures differentiated with this protocol (Figure 1B).

We concluded that the simultaneous combination of GMP-grade reagents (Table S2), standardised EB formation and regent-based rosettes isolation provides an efficient system for the derivation of lt-NES using a GMP-compatible platform.

### Neural differentiation potential of clinical grade hESCs lines

With a new GMP-compatible lt-NES derivation protocol developed we next examined its robustness by screening a panel of 6 clinically derived hESC (MasterShef) lines. MasterShef-3, −4, −7, −8, −10 and −11 were derived at, and obtained from, the Centre for Stem Cell Biology, University of Sheffield, under a HTA licence for potential clinical application (22510). With this approach, we aimed to also examine the specific neural differentiation potential of these clinical-grade lines which have been deposited in the UK Stem Cell Bank. Since for the derivation of lt-NES it is essential to generate neural rosettes, we decided to assess the differentiation potential based on the ability of the lines to give rise to morphologically distinct neural rosettes. The same number of cells for each line were differentiated into neural rosettes following our protocol and rosette formation was recorded by imaging the whole cell culture vessel with high definition imaging using BiostationCT twice daily for 5 days following EB plating (Figure 2A). Four out of six MasterShef lines (8, 4, 11 and 7), as well as an additional hESC research line from a different source, H9, were able to differentiate into neural rosettes, showing that the protocol is robust across several different lines (Figure 2A). Next, we scored each line for the percentage of neural induction by counting the numbers of EBs hosting neural rosettes in the whole-vessel images at the end of the induction, normalized to the total number of EBs attached (Figure 2B). The efficiency of neural induction is summarised in Figure 2C. We defined ‘good’ scores when more than 50% of the EBs carried rosettes, ‘medium’ scores when the value was below 50% but above 5%, and null when rosettes were undetected. Good neural induction scores were obtained regardless of the general level of spontaneous differentiation in pluripotency maintenance conditions assessed by daily morphological monitoring of the cultures for signs of differentiation (Figure 2C). Overall, these data demonstrate that our GMP-compatible protocol is suitable for an efficient neural differentiation of several clinically relevant hESCs without the need for cell-line specific optimisation.

**Figure 2.**
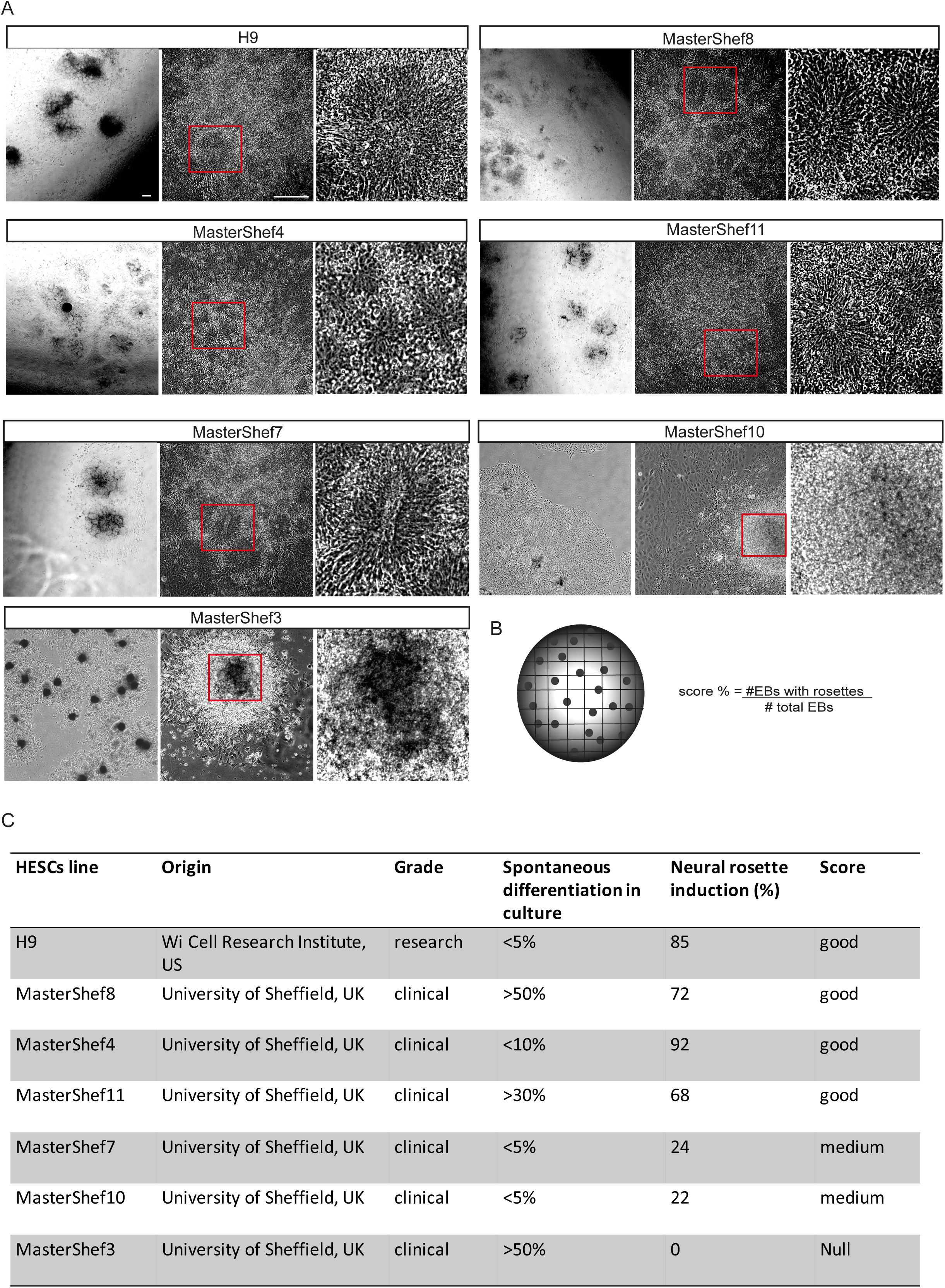
Neural differentiation potential of clinical grade hESCs lines. A) Representative phase contrast images of H9 and clinical-grade hESCs differentiated under GMP-compliant protocol into neural rosettes. Enlarged neural rosettes are visible in the right-hand side panels. Scale bars 200 μm B) Formula for the calculation of neural induction efficiency based on neural rosette in hESCs lines. C) Summary of screening of research and clinical-grade hESCs for neural induction capacity under GMP-compatible protocol.

### Establishment and characterization of GMP-compatible lt-NES

After screening clinical grade hESCs with our protocol, we examined if bona-fide GMP-compatible lt-NES could be derived and maintained from these lines. We successfully established new lt-NES from both a good score, MasterShef 8 and a medium score line, MasterShef 7, hESC line, which we named NES8 and NES7, respectively. NES7 and NES8 showed typical lt-NES morphology and self-organized in rosette-like clusters (Figure 3A), similarly to the published research-grade lt-NES AF22 (Figure S3). Furthermore, they homogeneously expressed lt-NES markers Nestin, SOX2, DACH1, PLZF and the polarity marker ZO-1 by immunofluorescence (Figure 3A). Consistently, NES7 and NES8 also expressed high level of lt-NES specific markers by Q-PCR, comparably to control lt-NES AF22 (Figure 3B). Our cells also preserved particularly useful features of lt-NES in the context of cell therapy manufacturing, such as good recovery after freeze-thaw (Figure 3C) and exponential proliferation in GMP conditions (Figure 3D) while retaining a rosette-like morphology (Movie 2). Finally, NES grown on laminin 521 maintained a normal karyotype (results from 30 spreads) after more than 15 passages (Figure 3E).

**Figure 3.**
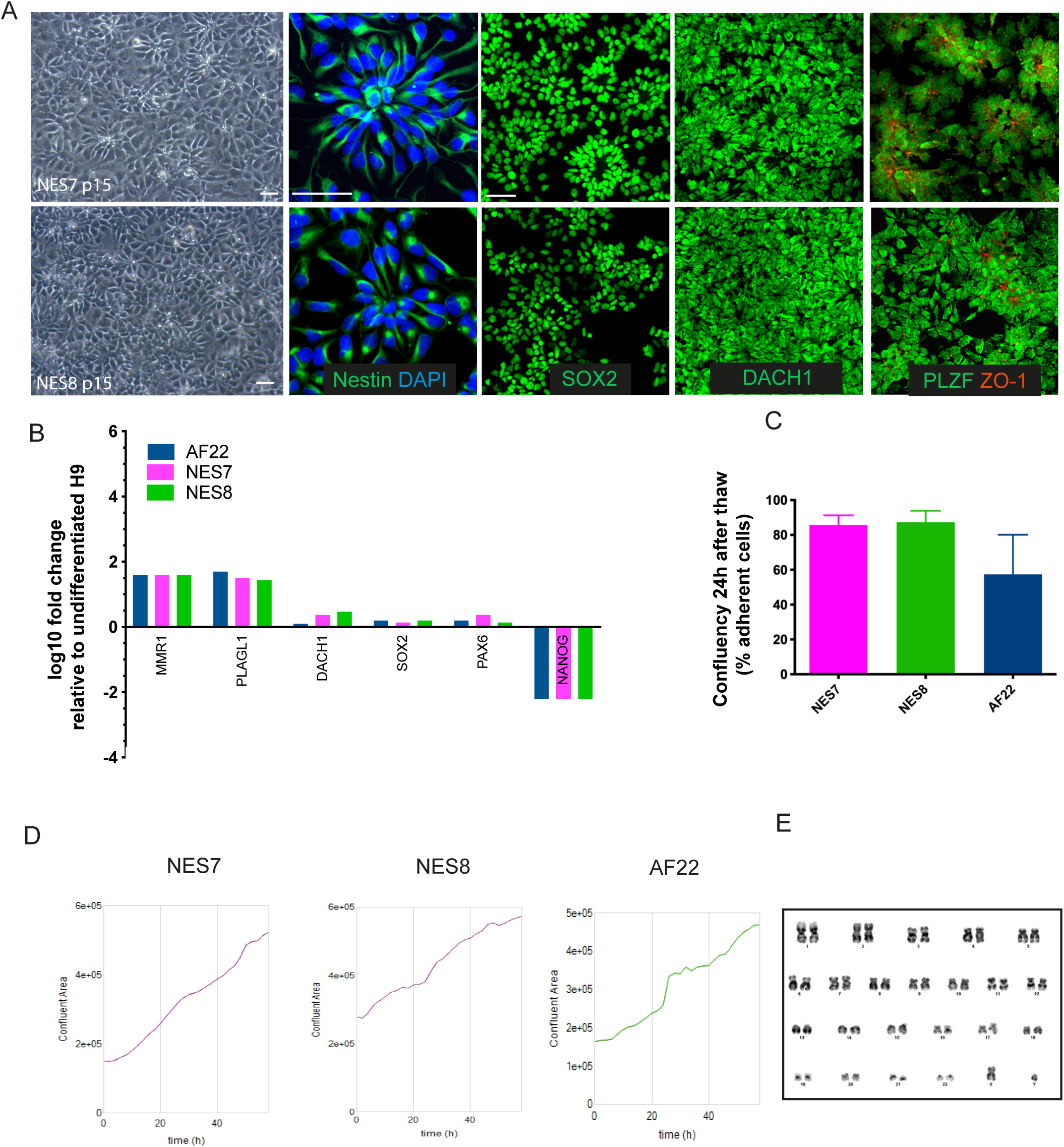
Establishment and characterization of GMP-compatible lt-NES. A) Representative phase contrast showing morphology of lt-NES derived from clinical grade MasterShef 7 (NES7) and 8 (NES8). Scale bars 20 μm. NES7 and NES8 cells were immunostained for the lt-NES markers Nestin, SOX2, Dach1, PLZF and the polarity marker ZO-1. Scale bars 50 μm B) Gene expression levels of lt-NES markers in NES7 and NES8 compared to research-grade AF22 lt-NES line and assessed by Q-PCR. (n=3). C) Recovery after thaw of NES cells at 24h. Graph shown as mean plus SEM (n=3). D) Representative growth curve of NES7, NES8 and control AF22 cultured under GMP-compatible conditions on L521. E) Normal karyology of established NES7 at passage 15 cultured in GMP-compatible system.

These results demonstrate that GMP-compatible lt-NES derived from clinical-grade hESCs are comparable to research grade lt-NES in morphology, markers and proliferative attributes.

### Spontaneous and directed GMP-compatible differentiation of lt-NES

Lt-NES have been shown to have a spontaneous bias towards hindbrain phenotypes, nevertheless retain multipotency and can be directed to differentiate into other neuronal cell types (Lundin et al. 2018, Falk. et al. 2012, Koch et al., 2009). Therefore, we examined whether our GMP-compatible lt-NES were able to differentiate into neurons in GMP-compatible differentiation conditions, using laminin 521 as substrate.

Firstly, we assessed our lt-NES spontaneous neurogenic potential by removing growth factors and allowing differentiation in neurogenic GMP media for 21 days by adapting research grade protocols (Falk et al. 2012). We observed that both NES7 and NES8 had a high neurogenic potential, giving rise to a homogenous and interlinked network of neurons over the course of the differentiation (Movie 3 and Figure 4). Indeed, the differentiated lt-NES expressed the neuronal marker Beta III tubulin (Figure 4A). Moreover, we confirmed the GABAergic propensity of the lt-NES since a large number of neurons were positive for markers GABA, in line with previous reports (Tailor et al. 2013) and similarly to the control line AF22 (Figure 4A).

**Figure 4.**
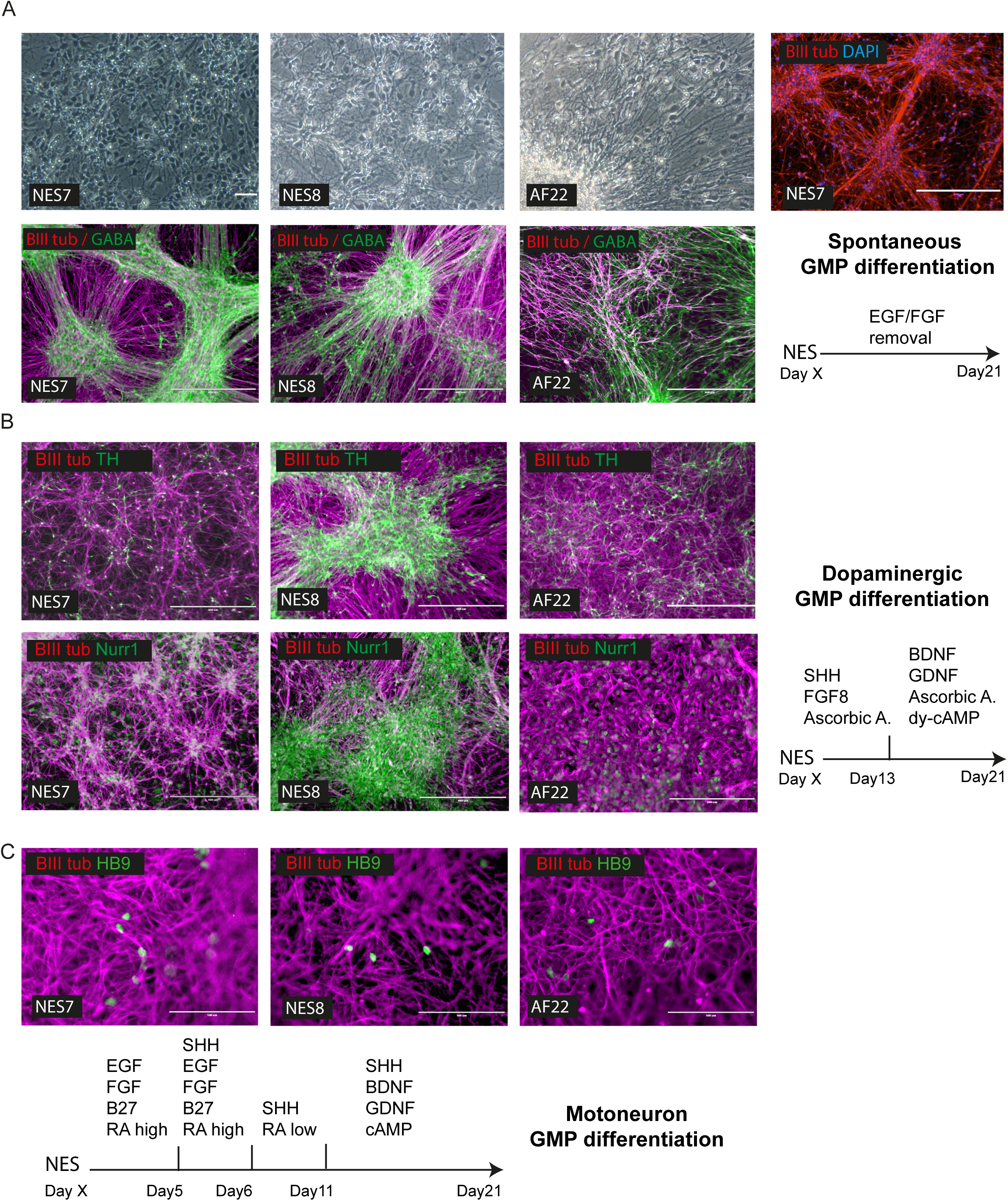
Spontaneous and directed GMP-compatible differentiation of lt-NES. A) Spontaneous differentiation of NES lines into neurons under GMP-compatible conditions. Spontaneous neurogenic potential shown by phase contrast (Scale bars 100 μm) and immunostaining for neuronal marker BIII Tubulin (NES7; scale bar 400 μm). Typical hindbrain bias of lt-NES shown by expression of GABAergic markers GABA (Scale bars 400 μm). B) Directed differentiation of NES lines into midbrain dopaminergic neurons under GMP-compatible conditions. Immunostaining images showing positivity for dopaminergic markers tyrosine hydroxylase (TH) and Nurr1. Scale bars 400 μm (200 μm for AF22 Nurr1). C) Directed differentiation of NES lines into motoneurons under GMP-compatible conditions showing motor neuron marker HB9 expression by immunofluorescence. Scale bars 100 μm.

Secondly, we examined the potential of our lt-NES to be redirected to alternative neuronal types under GMP differentiation conditions. Substituting reagents from previous lt-NES research methods (Koch et al., 2009) with GMP-grade equivalents, we tested differentiation towards a dopaminergic phenotype, which could have interest for cell transplant studies for Parkinson’s disease. Our results show that our GMP lt-NES method was able to give rise to neurons expressing dopaminergic markers Tyrosine Hydroxylase (TH) and Nurr1 after 21 days (Figure 4B).

Finally, lt-NES could also be directed towards a motoneuron phenotype in GMP conditions recapitulating research-grade protocols (Koch et al., 2009), even if much less efficiently than for the dopaminergic differentiation, as showed by detection of the mature motoneuron marker HB9 by immunofluorescence (Fig 4C).

In conclusion, our data demonstrated the feasibility of a fully defined and GMP-compatible protocol for the derivation and differentiation of hESCs into neurons via stable and expandable intermediate progenitors lt-NES.

## DISCUSSION

In this study, we established a system for the derivation, maintenance and differentiation of neuroepithelial stem cells lt-NES under GMP-compliant conditions suitable for clinical applications. We substituted each reagent of the original research-grade protocols (Falk et al., 2012; Koch et al., 2009) with available reagents of sufficient quality standards to allow their clinical application, so called GMP-approved reagents or cell-therapy-grade reagents. Manufacturers of such reagents have either lodged “drug master files” with regulatory authorities or are able to provide the detailed quality documentation required to perform a full risk assessment, which would in turn satisfy an appropriate regulator. In a few circumstances, when such grade was not readily available, we implemented reagents that are fully defined or which are in the process of achieving this standard by the manufacturers.

We also optimized the original differentiation protocol for transition to cell manufacturing environments. We found that a controlled and standardized EB formation was not only desirable but necessary for use of GMP Essential 6 media in the first phase of the protocol. In practice this led to higher reproducibility and yield of neural induction compared to the traditional KSR system. It would be interesting to investigate across a greater number of differentiation platforms whether the use of GMP-grade reagents provides an avenue for improvement of current research protocols. It would also be interesting to evaluate whether the choice of routine pluripotency maintenance conditions affects downstream differentiation.

A critical aspect of hESCs is their intrinsic line-to-line variability (Cahan and Daley 2013). In the context of cell therapy, our new protocol demonstrated robustness when applied to six different clinical-grade hESCs lines with a 66.6% efficiency. Moreover, lt-NES derived from hESCs with different neural differentiation propensities expressed similar characterization attributes between each other and to a control research-grade line. These findings confirm previous data reporting that lt-NES derivation may circumvent upstream differences between hPSC lines (Falk. et al., 2012), but here also showing comparability across different derivation conditions (i.e. GMP vs research protocol). As expected, a few of the screened lines were not able to differentiate efficiently under this protocol, reflecting the known characteristic of hESCs lines to have different developmental potentials. In this regard, this study strengthens the view that screening a panel of pluripotent lines is crucial for cell therapy applications.

We also established that GMP-grade maintenance conditions support the features of self-renewing lt-NES: rosette-like morphology, homogeneous and stable expression of neural rosettes markers, long-term expansion in EGF/FGF, resistance to repeated freeze/thawing and stable karyotype. Moreover, our lt-NES displayed high neurogenic potential towards GABAergic subtypes upon growth factors removal, in line with their hindbrain identity (Koch et al., 2009; Falk et al., 2012). lt-NES could be successfully re-specified towards adjacent regional cell types such as dopaminergic neurons and to a less degree also to motor neurons under GMP-culture conditions, confirming their multipotentiality. Therefore, with these protocols we intend to offer a flexible starting point and cut the burden of time-consuming and expensive process development.

The clinical future of pluripotent-stem cell-derived therapeutics will likely depend on our ability to tackle several roadblocks associated with the development of cell manufacturing processes, of which adaptation to suitable qualified materials is a crucial phase (Williams et al. 2016; Whiting et al., 2015; Ratcliffe et al., 2013; Abbasalizadeh et al., 2013). This study shows that translation of research-grade protocols to GMP-compliant protocols can be effectively achieved, and that making these changes can lead to robust and efficient processes. Stem cell researchers looking at transitioning to clinic can take our results as positive evidence that the original research protocol blueprint can be maintained and built upon. However, the time and cost commitment to achieving this should not be under-estimated, and with few reports in the literature to document these processes many developers have to start from the beginning.

Overall, the findings of the present report demonstrate feasibility of a GMP-compliant differentiation protocol for intermediate neural progenitors that are easy to expand, bankable and amenable to downstream differentiation into different neuronal sub-types. In the context of the cell therapy field, we report pre-screening of the neuronal differentiation capacity of 6 clinical-grade MasterShef hESC lines deposited in the UK Stem Cell Bank and established a resource in GMP lt-NES that could be used for further optimization depending on the required therapeutic goal. Recently, a method for differentiating lt-NES towards astroglia has been developed, opening new avenues for the downstream therapeutic use of GMP lt-NES (Lundin et al., 2018). Given its robustness and flexibility, our protocol could be applied for the generation of GMP-compliant neural progenitors that are potentially employable for a variety of neurological therapies, for cell manufacturing scalability studies, drug screenings and other biomedical research applications (Huang et al., 2019).

## MATERIALS AND METHODS

### Cell lines and culture methods

Derivation of the MasterShef-3, −4, −7, −8, −10 and −11 cell lines was performed in the Stem Cell Derivation Facility at the Centre for Stem Cell Biology, University of Sheffield, under HFEA licence R0115-8-A (Centre 0191) and HTA licence 22510, in a clean room setting, following strict standard operating procedures. The embryos used to derive MasterShef-3, −4, −7, 10 (frozen embryos) and MasterShef-8 and −11 (fresh embryos) were donated from different Assisted Conception Units, following fully informed consent, with no financial benefit to the donors, and were surplus or unsuitable for their IVF treatment. Briefly, the embryos, were cultured using standard IVF culture media (Medicult), to the blastocyst stage. Following removal of the trophectoderm using a dissection laser the embryos were explanted whole onto either mitotically inactivated human neonatal fibroblasts (human feeders) in the case of MasterShef-3, −4, −7, 8 and −10 or onto Laminin-511 (Biolamina) in the case of MasterShef11. Derivation media for MasterShef-3, −4, −7 and −8 was standard KSR/KODMEM (Life Technologies) medium whilst MasterShef-10 and −11 were derived in Nutristem medium (Biological Industries). All cell lines were initially maintained at 37°C under 5% O_2_/5% CO_2_, until the lines were established, after which maintenance switched 5% CO_2_ in air at 37^°^C. Cultures were passaged using a manual technique, cutting selected colonies under a dissection microscope at an average split ratio of 1:2 every 7 days. All cell lines have been deposited at the UK Stem Cell bank (https://www.nibsc.org/ukstemcellbank). The H9 cell line was obtained from the WiCell Institute, US.

HESCs were routinely maintained on recombinant VTN-N Vitronectin (A14700, Life Technologies), also tested on the prototype CTS™ (Cell Therapy Systems) Vitronectin with similar results (now A27940, Life Technologies), and GMP Essential 8 (A1517001, Life Technologies). For routine passaging, cells were washed once with CTS™ DPBS−/− (A1285601, Life Technologies) and incubated at room temperature for 1-2 minutes with GMP EDTA (15575020, Invitrogen). After aspiration of EDTA, colonies were gently detached as small clumps and passaged at a ratio of 1:6 without centrifugation. Cells were frozen in animal-free freezing medium, CryoStem (K1-0640, Geneflow), and thawed in presence of GMP ROCK inhibitor Revitacell (1:100, A2644501, Life Technologies). For traditional research-grade differentiation and reagents, refer to the supplemental experimental procedures.

### Establishment of GMP lt-NES

Undifferentiated hESCs were dissociated into single cells with StemPro Accutase* (A1110501, Life Technologies) for 2-3 minutes at 37oC, suspended into GMP Essential 6 (A1516401, Life Technologies), counted with a haemocytometer and centrifuged at 300g for 5 minutes. Cells were suspended at a concentration of 3 × 10^6^ into 1.5ml of (E6) plus Revitacell, and gently mixed into one well of Aggrewell 800 (Stem Cell Technologies) previously centrifuged at 2000g with 500 μl of E6 plus Revitacell. EBs formed after 24 hr and the media was carefully and completely replaced with fresh E6. From day 2 till day 4 EBs were fed daily with half media change within the Aggrewell. At day 5, EBs were detached from the Aggrewell using a p1000 tip while a large bore tip (Starlab, E1011-9618) was used for careful collection and deposition of the EBs on the top of a 37 μm reversible strainer (Stem Cell Technologies). Multiple cycles were performed with E6 until all the EBs were removed. EBs were then plated onto 1 well of a 6-well plate (Corning) coated overnight with 10 μg/ml of xeno-free human recombinant Laminin 521 (LN521, Biolamina) prepared in GMP DPBS^+/+^ (A1285801, Life Technologies) by reversing the strainer and washing the EBs into the plate with GMP N2 media (CTS™ DMEM-F12, A1370801; CTS™ N2 1:100, A1370701; CTS™ B27 1:1000, A1486701; 1% GMP Glutamax, A12860-01; Life Technologies). Neural induction was induced for 3-5 days by changing GMP N2 media daily. Neural rosettes were derived between day 3 to day 5 by addition of STEMdiffTM Neural Rosette Selection Reagent (05832, Stem Cell Technologies) for 45 mins-1 hr at 37oC. The rosettes were gently detached with N2 media directed with a p1000 tip on the visible rosette clusters. Purity of selection was checked under the microscope for detachment of rosette clusters and non-differentiated cells. Removed rosettes were collected in a tube and new media was used to continue selection until 70% of the rosettes were collected. Rosettes were centrifuged at 300g for 5 minutes and suspended into 400 μl of N2 media plus 10 ng/ml of GMP FGF and GMP EGF (233- GMP-025, 236-GMP-01M; Bio-techne), named N2 EF media, plus Revitacell. Cells were plated into one to 4 wells of a 48-well plated pre-coated with 10 μg/ml Laminin 521 avoiding over pipetting and formation of single cells. Critical steps and troubleshooting: Successful derivation of lt-NES depends on proper attachment. It is recommended to prepare several laminin plates of various sized during early derivation in order to have reserve plates readily available in the event that attachment is not optimal. A high cell density is required for lt-NES to survive and proliferate, around 70%.

Cells were fed daily with GMP N2 EF media. Once confluent, lt-NES were dissociated with Accutase for 1 min at 37oC, collected by pipetting on the surface and suspended into 10 ml N2 media prior to centrifugation at 300g for 5 mins. Cells were passaged at a split ratio of 1:1 from a 48 well format to a 6 well plate with addition of Revitacell for the first 24 hr. Once cells were in 6-well format, Revitacell was not used during passaging and lt-NES were split at a ratio of 1:2 or 1:3.

### GMP lt-NES maintenance

lt-NES were routinely cultured in GMP N2 EF media on 10 μg/ml Laminin 521. Cells were split every 3-4 days when sub confluent with incubation with Accutase 1-2 mins at 37°C (without waiting for the cells to be floating in the media) and suspended into 10 ml N2 media before centrifugation at 300g for 5 mins. Cells were frozen with Cryostem. lt-NES were thawed at 37°C for 2 minutes and immediately resuspended into 10 ml N2 media, centrifuged at 300g for 5 minutes and plated in N2 EF media plus Revitacell for the first 24 hr.

### Spontaneous differentiation of lt-NES

lt-NES were plated at a density of 40,000 cells/cm^2^ on Laminin 521 coated plates in N2 media plus Revitacell for 24hr. The next day, media was changed to terminal differentiation media composed of 50:50 parts of CTS™ DMEM-F12 (with CTS™ N2 1:100) and CTS™ Neurobasal (A1371201, Life Technologies) (with CTS™ B27 1:50) media plus 300 ng/ml cAMP (Sigma Aldrich). Spontaneous differentiation was induced with the media above for 21 continuous days.

### Directed GMP differentiation into dopaminergic neurons

lt-NES were plated at a density of 40,000 cells/cm^2^ on Laminin 521 coated plates in N2 media plus Revitacell for 24hr. The next day, media was changed to dopaminergic patterning medium composed of CTS™ DMEM-F12 (with CTS™ N2 1:100) plus freshly added 200 ng/ml GMP Sonic Hedgehog (SHH, 130-095-727, Miltenyi Biotec), 100 ng/ml GMP FGF-8b (130-095-740, Miltenyi Biotec) and 160 μM Ascorbic Acid (95210-250G, Sigma Aldrich). Cells were cultured in dopaminergic patterning medium for 2 weeks. On day 14, media was changed into terminal differentiation medium composed of equal parts of CTS™ DMEM-F12 (CTS™ N2 1:100): CTS™ Neurobasal (CTS™ B27 1:50) plus 20ng/ml GMP BDNF (248-GMP-025, Bio-techne), 10 ng/ml GMP GDNF (212-GMP-050, Bio-techne), 160 M Ascorbic Acid (Sigma Aldrich), 500 μM dy-cAMP (Sigma Aldrich). Cells were continuously fed with terminal differentiation media until day 21, when neurons are ready for immunofluorescence characterization.

### Directed GMP differentiation into motoneurons

lt-NES were plated at a density of 40000 cells/cm^2^ on Laminin 521 coated plates in N2 media plus Revitacell for 24hr. The next day, media was changed to motoneuron patterning medium composed of CTS™ DMEM F12 (With CTS™ N2 1:100, CTS™ B17 1:50) plus 10ng/ml GMP EGF (Bio-techne), 10ng/ml GMP FGF (Bio-techne) and 1 M Retinoid Acid (Sigma Aldrich). On day 5, the above media was supplemented with 1μg/ml GMP SHH (Bio-techne). From day 7, the concentration of Retinoid Acid was reduced down to 0,01 μM and EGF and FGF were completely removed. On day 12, media was changed to terminal differentiation media composed of equal parts of CTS™ DMEM-F12 (CTS™ N2 1:100): CTS™ Neurobasal (CTS™ B27 1:50) plus 20 ng/ml GMP BDNF (Bio-techne), 20 ng/ml GMP GDNF (Bio-techne), 50ng/ml SHH (Bio-techne) and 300 ng/ml cAMP (Sigma Aldrich).

## SUPPLEMENTAL INFORMATION

Supplemental information includes 1 figure, 3 movies and supplemental experimental procedures.

## AUTHORS CONTRIBUTION

L.V. performed, analysed the experiments and wrote the manuscript. C.D. assisted with maintenance of MasterShef lines cultures and experiments. Z.H. and H.M derived the clinical-grade MasterShef hESC lines and A.S. provided lt-NES AF22 line. A.S. and L.V supervised the study. All authors reviewed and assisted on the manuscript.

## ACKNOWLEDGEMENTS

This work was supported with a UK Regenerative Medicine Platform grant funded by the Medical Research Council, the Biotechnology and Biological Sciences Research Council and the Engineering and Physical Sciences Research Council. We thank Duncan Baker for cytogenetic analysis at Sheffield’s NHS Children Foundation Trust. We thank Dr. Anna Falk and Dr. Jignesh Tailor for precious advice on the research-grade protocols. We also would like to thank the late Dr. Nicholas Blair for his unconditional support and kindness.

