## Supplemental information for "GMP-grade neural progenitor derivation and differentiation from clinical-grade human embryonic stem cells"

### SUPPLEMENTAL FIGURES

#### FIGURE S1

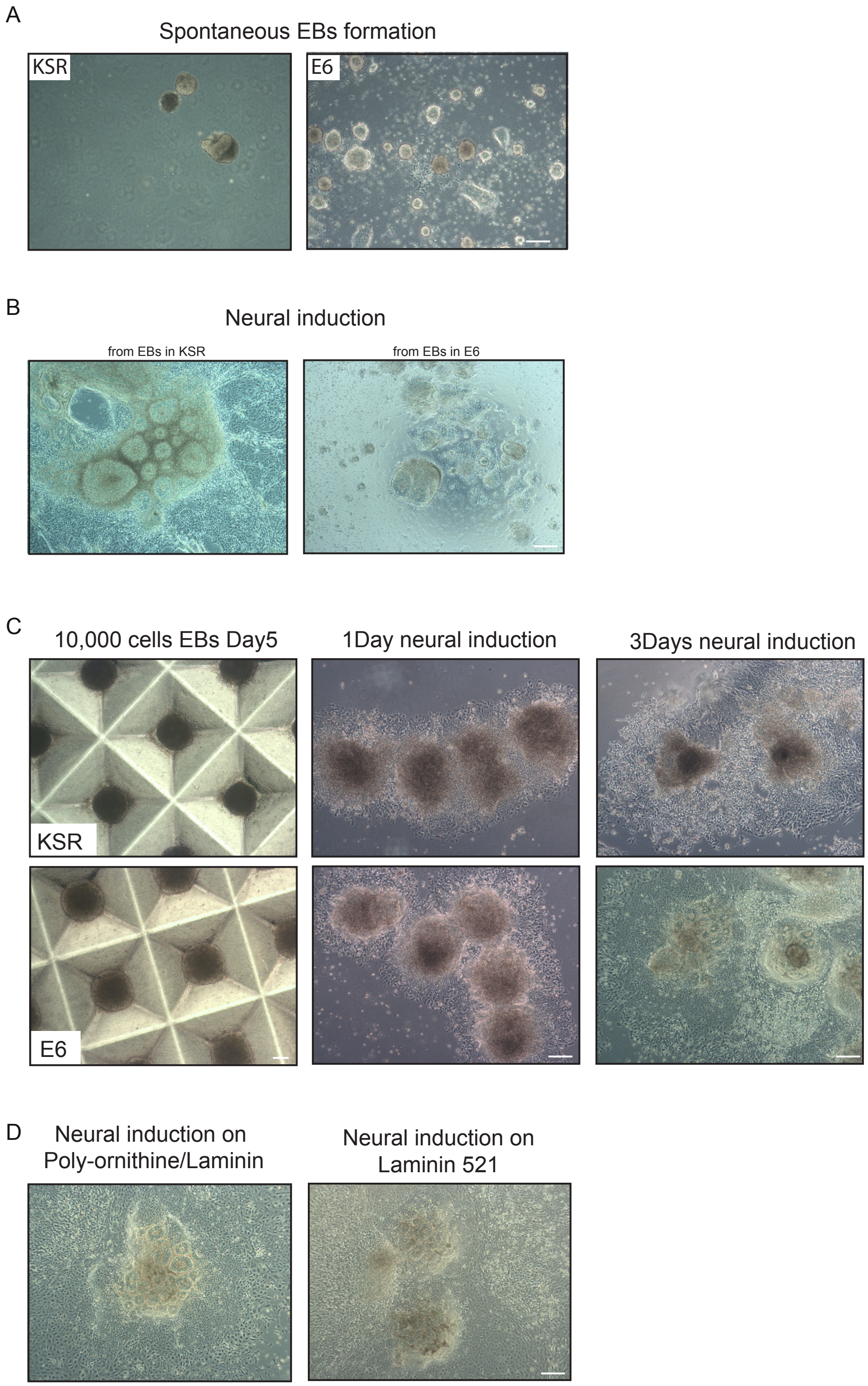

**Figure S1 Development of an efficient GMP-compatible protocol for lt-NES (Related to Figure 1)**

A) Representative phase images of embryoid bodies spontaneously formed from hESCs (H9) in suspension on low-adherence plates in KSR or Essential 6 media.
B) Representative phase images of neural rosettes induction from spontaneously formed EBs (from H9) in KSR versus Essential 6 media.
C) Representative comparison of standardized neural rosette induction (from MShef10) under research-grade KSR or GMP Essential 6 media.
D) Representative phase images of neural rosettes formation (from H9) on research-grade substrate made of poly-L-Ornithine/laminin versus the defined recombinant laminin 521 matrix.

Scale bars 100 $\mu$m

### TABLE S2

| Stage | **Research**<br>A combination of protocols from Koch et al. 2009; Falk et al. 2012; Tailor et al. 2013 | | **GMP**<br>**CTS= Cell therapy systems** *(serum-free, xeno-free and animal origin-free. Cell and gene therapy specific intended use statements)*<br>**GMP proteins** *are animal-free and cGMP compliant and recommended for clinical use* | | |
|---|---|---|---|---|---|
| | Reagent | Source | Reagent | Source | Identifier |
| Pluripotency | Irradiated Mouse Embryonic Fibroblasts | n/a | VTN-N Vitronectin | Life Tech. | A14700<br>now A27940 (CTS™) |
| | Stem cell media:<br><br>DMEM-F12<br>NEAA<br>L-Glutamine<br>KSR<br>$\beta$ –mercapto. | In house<br><br>Reagents from Invitrogen | Essential 8 | Life Tech. | A1517001<br><br>now A2656101 (CTS™) |
| | FGF | Peprotech | | | |
| | DPBS$^{-/-}$ | Life Tech. | CTS™ DPBS$^{-/-}$ | Life Tech. | A1285601 |
| | Collagenase | Invitrogen | EDTA | Invitrogen | 15575020 |
| | Stem cell media + 10 % DMSO | In house | CryoStem | Geneflow | K1-0640 |
| | | | Revitacell (Rocki) | Life Tech. | A2644501 |
| EBs | EDTA | Life Tech. | StemPro Accutase* | Life Tech. | A1110501 |
| | KSR:<br>Advanced DMEM-F12<br>KOSR<br>L-glutamine<br>NEAA | In house<br><br>Reagents from Life Tech. | Essential 6 | Life Tech. | A1516401 |
| | $\beta$ –mercap. | Sigma | | | |
| | | | Revitacell | Life Tech. | A2644501 |
| Rosette Induction | Poly-L-Ornithine | Sigma | Laminin 521 | Biolamina | LN521<br>(now also Cell therapy grade) |
| | Laminin (L2020) | Sigma | | | |
| | DPBS$^{-/-}$ | Life Tech. | CTS™ DPBS$^{-/-}$ | Life Tech. | A1285601 |
| | N2 media: | In house | CTS™ N2 media: | | |
| | DMEM F12 | Reagents from Life Tech. | CTS™ DMEM F12 | Life Tech. | A1370801 |
| | N2 (1:100) | | CTS™ N2 | Life Tech. | A1370701 |
| | B27 (1:1000) | | CTS™ B27 | Life Tech. | A1486701 |
| | Glutamine | | CTS™ GlutaMAX | Life Tech. | A12860-01 |
| Rosette selection | Manual Picking | n/a | STEMdiff™ Neural Rosette Selection (enzyme-free)* | Stem Cell Tech. | 05832 |
| | N2 media | As above | CTS™ N2 media | As above | |
| | FGF | R&D | FGF | Bio-Techne | 233-GMP-025 |
| | EGF | R&D | EGF | Bio-Techne | 236-GMP-01M |
| | | | Revitacell | Life Tech. | A2644501 |
| lt-NES expansion | N2 media + FGF and EGF | As above | CTS™ N2 media + FGF and EGF | As above | |
| | Poly-L-Ornithine<br><br>Laminin L2020 | Sigma | Laminin 521 | Biolamina | LN521 |
| | Trypsin | Invitrogen | StemPro Accutase | Life Tech. | A1110501 |
| lt-NES freezing | Stem cell media + DMSO | In house | CryoStem | Geneflow | K1-0640 |

*Defined reagents of which risk assessment will have to be conducted. Not cGMP at time of study

**FIGURE S3**

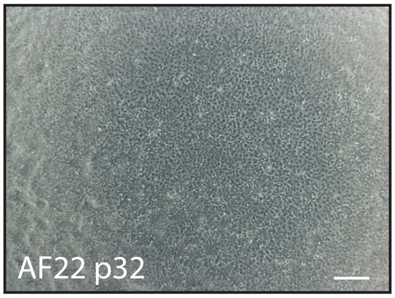


**Figure S3 Establishment and characterization of GMP-compatible lt-NES**

**(Related to Figure 3)**

A) Representative phase image of the established research-grade lt-NES line AF22. Scale bar 100 μm

#### SUPPLEMENTAL EXPERIMENTAL PROCEDURES

##### Traditional EB differentiation and research-grade reagents

Spontaneous EBs formations was achieved by detaching confluent hESCs with EDTA and directly depositing the cell aggregates at a split ratio of 1:1 in a ultralow-adherence plates in 10ml of research-grade KSR composed of Advances DMEM F12, 20% Knockout Serum Replacement, 1% Glutamine (all from Life Technologies) and 0.1mM β-Mercaptoethanol (Sigma). For comparison, GMP Essential 6 media (Life Technologies) was tested. Media was changed every other day for a total of 7 days by collection of the floating EBs to the bottom of a falcon tube for 10 mins followed by re-plating into fresch media. At day 7, EBs were seeded onto a plate prepared by coating with Poly-L-Ornithine solution (P4957-50ml, Sigma) for 30 mins at room temperature, washed twice with DPBS$^{-/-}$ (Life Technolgies) and then coated with 10 μg/ml of laminin (L2020, Sigma) overnight at 4$^{o}$C. Neuronal induction was induced in N2 base media as previously described (Falk et al. 2012).

##### Immunostaining

Cells were fixed for 10 mins with 4% parafolmaldehyde and then washes three times with PBS. For blocking and permeabilization, cells were incubated for 30 mins in PBS plus 10% donkey serum (Serotec) and 0.1% Triton X-100 (Sigma Aldrich). Cells were incubated overnight at 4$^{o}$C with primary antibodies diluted in PBS plus 1% donkey serum and 0.1% Triton X-100. Cells were then washed in 3 times in PBS and incubated for 1hr with fluorescent secondary antibodies (Life Technologies) in 1% donkey serum and 0.1% Triton X-100 (in PBS). Cells were washes 3 times with PBS and counter stained with DAPI (New England Biolabs). Cells were imaged with confocal LSM 700 microscope (Zeiss) and EVOS™ FL microscope (Life Technologies).

Antibodies were against: SOX2 (Bio-techne), Nestin (Abcam), DACH1 (Proteintech), PLZF (Life Technologies), ZO-1 (Bio-techne), GABA (Sigma), TUJ1 ( Biolegend), Tyrosine Hydroilase (Millipore), Nurr1 (Santa Cruz), HB9 (Insight Biotechnology).

**RNA isolation and Q-PCR**

RNA was extracted using the RNeasy kit following manufacturer instructions (Qiagen). An amount of 250 ng of RNA was retro transcribed with the RXN MAXIMA $1^{ST}$ strand cDNA synthesis kit (Fisher scientific) and diluted 1:40 in ultrapure water. Q-PCR assay was performed with FG, TAQMAN Gex Master Mix (Life Technologies) in a 20 μl reaction including 5 μl of RNA and 1μl of TAQMAN GENE EX Assays. TaqMan assays used were: SOX2 (Hs01053049_s1), PAX6 (Hs00240871_m1), DACH1 (Hs00362088_m1), PLAGL1 (Hs00414677_m1), MMR1 (Hs00201182_m1), NANOG (Hs04260366_g1), PBGD (Hs00609296_g1). Q-PCR was run and analysed with a QuantStudio 12K flex real time machine (Life Technologies) following the TAQMAN comparative program. Samples were normalized to housekeeping gene PBGD.

**Karyology**

NES were prepared for karyotype analysis following methaphase arrest by incubation with 2 μg/ml Colcemid for 4 hr at 37°C. Cells were then dissociated in 0.25% trypsin/EDTA (Stem Cell Technologies) and then incubated with hypotonic solution of

0.00375M KCl for 10 mins at room temperature. Cells were centrifuged at 100g for 8 minutes and the pellet suspended drop by drop in fixative solution (3 parts Methanol and 1 part acetic acid; Sigma). Centrifugation and fixation were repeat 3 times before sample were left in 0.5 ml fixative. Cells spreads were controlled prior karyotyping by dropping 10 μl of cell suspension onto a histology slide from a height of approximately 50 cm. Dried samples were stained with DAPI mounting medium (Sigma Aldrich) and harvest of chromosomes clusters confirmed under a fluorescent microscope. Karyotype analysis on fixed samples was performed by Sheffield Diagnostic Genetic Services (Sheffield Children's Hospital, UK).

**Biostation CT imaging**

Cells undergoing neural induction were imaged at a 2x and 10x magnification with Biostation CT (Nikon) to acquire full-well tiling images every 12 hr for 5 days starting from 1 hr after plating of EBs onto laminin. For proliferation analysis, lt-NES were passaged normally at a ratio of 1:3 into 6-well plates and imaged every 2 hr for 3 days with a 2x2 tiling, 20x magnification. Images were then analysed with CL Quant software following the cell proliferation recipe (Nikon). For terminal differentiation, lt-NES were plated at a density of 40,000 cells/cm$^2$ and imaged at 20x magnification every 12 hr for 21 days under spontaneous differentiation protocol. All movies were created with CL Quant (Nikon).
